# Bottleneck stages determine resilience in successional communities

**DOI:** 10.64898/2026.07.31.742051

**Authors:** Nasser Rabi

**Affiliations:** W. K. Kellogg Biological Station, Michigan State University, Hickory Corners, Michigan 49060; and Program in Ecology and Evolutionary Biology, Michigan State University, East Lansing, Michigan 48824; Department of Integrative Biology, Michigan State University, East Lansing, Michigan 48824

**Keywords:** Succcession, resilience, Markov chains

## Abstract

Successional communities often recover slowly because progression stalls at persistent stages that resist replacement. Here, we show that resilience in such systems is frequently governed by a single bottleneck stage with the lowest effective exit rate. Using empirical transition matrices from intertidal and plant communities, we demonstrate that altering the bottleneck has a much larger effect on resilience than modifying any other stage. We then show that this bottleneck principle emerges naturally from both Markov and continuous-time models of succession. Specifically that the slowest return to equilibrium is controlled primarily by the stage with the smallest effective exit rate, which also dominates the mean first-passage time to late succession. These results provide a simple biological interpretation of resilience in successional communities and suggest that management efforts are most effective when they target the stage that limits the pace of succession.

## Introduction

Many sessile communities exist in successional sequences characterized by continual replacement of competitors that is reset by disturbances. Early successional species are often strong colonists but weak competitors, while later-arriving species tend to persist longer, allowing succession to be modeled as a fairly predictable sequence in which fast colonists are gradually replaced by superior competitors (Cadotte et al. 2006; Turnbull et al. 1999). The dynamics of such a sequence determines the time until a specified community can rebound from disturbances, or its resilience (Halpern 1988; Johnstone et al. 2016).

In many systems, succession appears to slow or arrest at persistent stages whose occupants resist replacement and thereby delay the advancement of the community towards a mature state. Such recovery-limiting stages are common across successional systems. In old fields, competition with invasive species and poorer dispersal of native species can arrest development during succession (Cramer et al. 2008; Jakovac et al. 2021). In forests, persistent fern, bamboo, or shrub layers can trap disturbed stands in arrested succession by forming dense shade to exclude later stages (Royo and Carson 2006). Likewise, in marine systems algal turfs and long-lived sessile organisms delay succession towards later successional states (Connell and Slatyer 1977; Hill et al. 2004). Across these systems, succession is often stalled by a small subset of species, suggesting that a community’s path towards equilibrium may be constrained less by the full network of species replacements than by a subset of stages that locks communities in place. This raises a natural question: which stages determine the rate at which the equilibrium community is approached, and hence its resilience?

Here we show that the quantity governing resilience in successional systems often admits a simple biological interpretation. Specifically, that resilience in successional communities is often governed disproportionately by a single *bottleneck stage*: the species or patch type with the lowest effective exit rate, which sets the timescale of return to the equilibrium community. Using empirical transition matrices from intertidal and plant communities, we show that this bottleneck stage disproportionately governs resilience in real successional systems, and that altering properties of other species tends to have comparatively little impact on resilience. We then show that this bottleneck principle emerges naturally from classical models of succession: although succession is often treated as the aggregate outcome of many species transitions, resilience is often limited primarily by stages that are unlikely to be replaced. This principle makes the timescale of return to equilibrium both biologically interpretable and, in principle, more easily managed. Throughout this paper, equilibrium denotes the stationary distribution of patch types maintained under ongoing disturbance and succession, not the complete occupation of a particular “climax” species/stage.

### Bottlenecks in empirical systems

Empirical successional systems are rarely observed reaching equilibrium, making resilience difficult to estimate from time series alone. However, succession in natural communities can be modeled well by a Markov transition matrix *A*, where *A_ij_* denotes the probability species *i* is replaced by species *j*, and the diagonal elements represent self-replacement (i.e., persistence of the same successional state) (Horn 1975; Waggoner and Stephens 1970). In these models, each patch occupies a discrete successional state, and the entries of A describe the probabilities that one state replaces another between observation periods. When parameterized, one can compute an equilibrium stable distribution of patch types, as well as a variety of sensitivity and stability indicators (Caswell 2013). While models with continuous dynamics can capture more realistic processes, such as density-dependent replacement probabilities or variable patch conditions (Amarasekare and Possingham 2001; Miller and Allesina 2021), discrete-time Markov models have successfully predicted successional dynamics in a variety of natural communities and provide a useful, analytically tractable framework for studying resilience.

Here we examine resilience in two empirical Markov transition matrices from sessile communities in which Markov chains were shown to predict stationary community composition well: a rocky intertidal assemblage from the Gulf of Maine (Hill et al. 2004) and a disturbed successional plant community from a sandy dry grassland (Baasch et al. 2010). Resilience is computed by the subdominant eigenvalue *λ*_2_ of the Markov transition matrix, which gives the rate of convergence towards a stationary distribution of patch types (Caswell 2019). In discrete Markov chains, smaller values of *λ*_2_ indicate faster return to equilibrium. Hence 1/*λ*_2_ can be interpreted as a metric of resilience.

To assess resilience in each empirical community, we conduct a simple perturbation analysis on the transition matrix. Each stage has its self-replacement (diagonal) probability reduced by a fixed amount. The removed probability was redistributed among forward successional transitions, effectively creating additional pathways by which a patch could leave that stage while preserving the total transition probability. The subdominant eigenvalue *λ*_2_ is then calculated to quantify the effect of this perturbation on resilience (Fig. 1). Because *λ*_2_ sets the timescale of the return to equilibrium, species whose perturbation produces the largest reduction in *λ*_2_ exert strong influence on community resilience.

**Figure 1:**
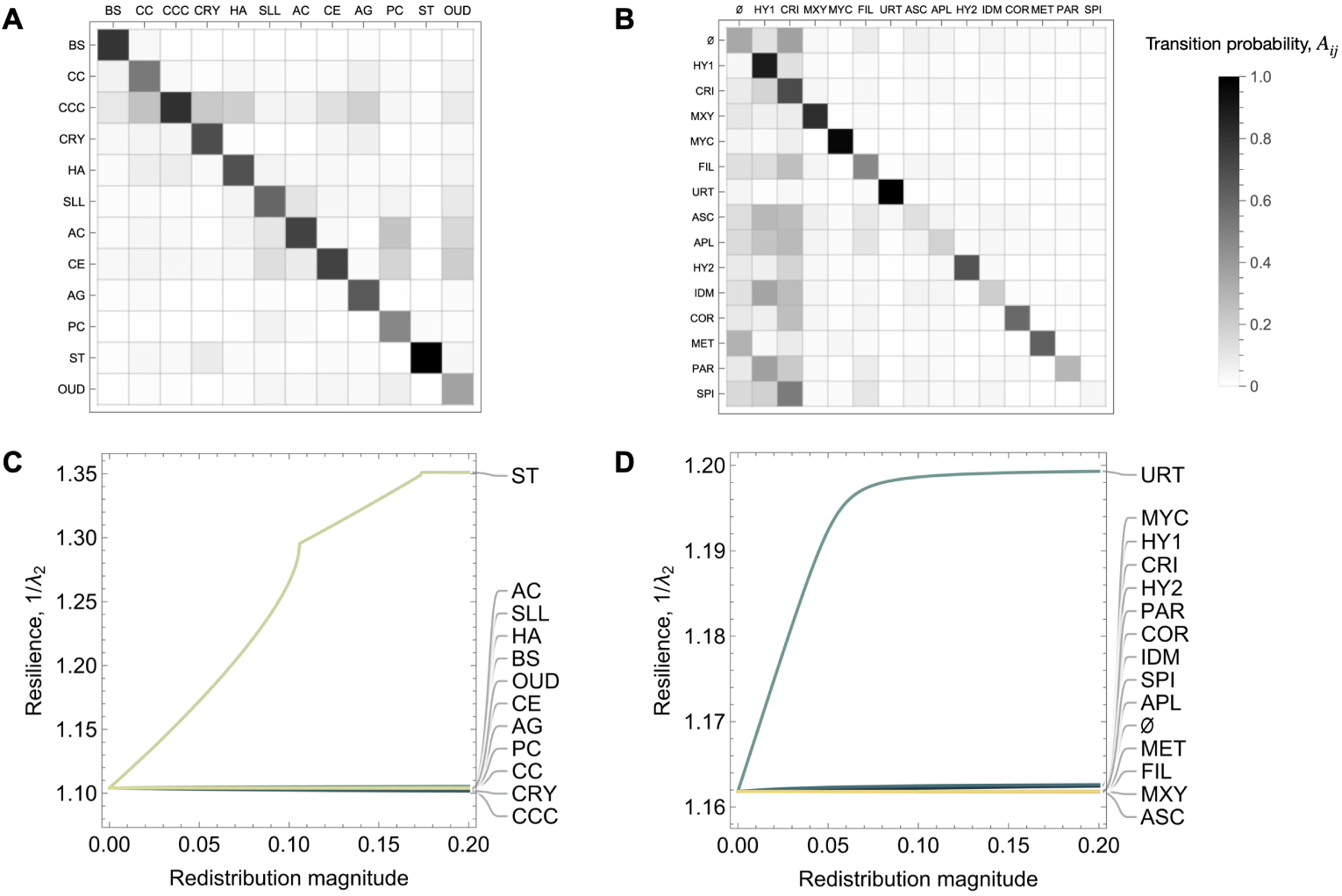
Empirical demonstration of bottleneck principle. Markov transition matrices of A) the successional intertidal community in Hill et al. (2004) and B) a sandy grassland plant community in Baasch et al. (2010). Panels (C), and (D), show the magnitude of the resulting subdominant eigenvalue (resilience of the entire community) when transition probabilities are redistributed from each diagonal element to the later stages in the successional community (specifically, OUD (“other/undefined communities”) in panel (C) and SPI (*Spirorbis spirorbis*) in panel (D).

Despite their biological differences, both communities show the same qualitative structure. Their transition matrices are strongly diagonally dominant (Fig. 1A,B), indicating that species self-replacements have the highest probability. In each system, one stage is markedly more persistent than all others: a sea anemone (*Urticina crassicornis*, URT) in the intertidal community and the “shrubs and trees” stage (ST) in the grassland system. These stages have the highest self-replacement probabilities in their respective communities (URT = 0.863; ST = 0.93), indicating unusually long residence times and identifying them as the dominant slow-turnover stages in each system.

In both systems, resilience is concentrated almost entirely in these bottleneck stages. Reducing self-replacement in the bottleneck produces a strong decrease in *λ*_2_ hence increasing community resilience, whereas redistributing transition probability from any other stage has little to no effect on resilience (Fig. 1C,D). This contrast is counterintuitive; although many transitions participate in succession, most do not limit the time to reach equilibrium because patches move through them relatively quickly. As a result, altering replacement rates elsewhere in the network does little to advance succession unless those changes shorten residence time in the bottleneck itself.

The consequent change in resilience depends on the severity of the bottleneck. Because ST has an exceptionally low *exit rate*, or probability of leaving that state, perturbing it produces a larger increase in resilience than an equivalent perturbation to URT. Across both communities, the same empirical pattern emerges: resilience is not limited diffusely by the full transition network, but disproportionately by one persistent stage that holds the community in place. Below, we show that this pattern emerges naturally from two widely used models of succession.

### Model and results

Theoretical models exist to study successional dynamics in seasonally fluctuating species (Klausmeier and Litchman 2012) and in spatially heterogeneous environments (Miller and Allesina 2021), but few have been successful at quantitatively predicting community composition. While modeling succession as a Markov process has proved useful across a variety of systems (Caspersen and Pacala 2001; Hill et al. 2002, 2004) its value as a theoretical framework for studying the mechanisms of succession has received comparatively little attention. Here, the theoretical goal is not to re-establish the result that *λ*_2_ governs resilience in Markov systems, but to derive a valuable biological interpretation for resilience.

In the Markov model of succession originally introduced in Horn (1975), each patch in the landscape occupies one of several discrete states, representing a species. Transitions among these states occur according to a fixed transition matrix A, which encodes the probabilities of replacement or disturbance between consecutive time steps. In the original formulation of Horn (1975), a patch begins in the empty or “disturbed” state and can be colonized by a competitively superior species with probability *e_i_*. Each species is affected by disturbance with probability *d_i_*, returning patches to the empty state so that even the most competitive (or “climax”) species is not absorbing. The resulting transition matrix, which moves the system from the vector of patch frequencies *p_t_* to *p_t_*_+1_, takes the form:

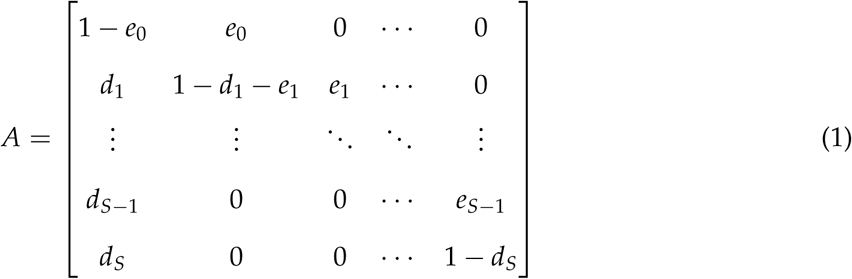

Here, the entry in row *i* and column *j* gives the probability that species *j* succeeds species *i*. For a system with *S* species, there are thus *S* + 1 possible states, including the disturbed or empty patch. This hierarchical structure in which disturbance resets succession and species advance along a sequence from poor competitors to superior competitors is common across a wide variety of systems, including forests ((Culver 1981; Korotkov et al. 2001)), intertidal communities (Hill et al. 2004) and nurse plant communities (McAuliffe 1988). Fundamentally the dynamics described in Eq. (1) describe a competition-colonization tradeoff, with early successional species possessing lower competitive ability and greater colonization ability, with late species being the opposite. Moreover, most successional events are concentrated about the diagonal as in the empirical matrices (Fig. 1A,B). The stationary distribution of the Markov chain, defined by the vector 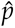 satisfying 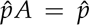, gives the long-term average of patches in each successional state. This equilibrium composition represents the frequency of each stage at long timescales.

While there are numerous ways to define resilience (Dakos and Kéfi 2022), here we define it as the rate until the system returns to equilibrium following a perturbation. In Markov chains, this is determined by the subdominant eigenvalues of *A*, which describe the rates at which deviations from equilibrium decay through time (Capdevila et al. 2020; Caswell 2013). Specifically, the magnitude of the subdominant eigenvalue determines the rate of return to equilibrium such that smaller magnitudes correspond to faster convergence, whereas values closer to one indicate slow return and hence lower resilience.

When species respond the same to disturbance (*d_i_* = *d*), the eigenvalues (except the dominant eigenvalue, 1) are given by *λ_i_* = 1 − *d* − *e_i_*. Since resilience is determined by the subdominant eigenvalue *λ*_2_, a greater value of *e_i_* corresponds to faster convergence of the community towards its steady state. Put another way, if a species is easily replaced (large *d* + *e_i_*), a patch can more easily move forward to later successional stages. Conversely, a slow rate of replacement (small *d* + *e_i_*) slows the rate equilibrium is attained.

Although it may seem that resilience should depend on the full network of transitions among species, in this model it is controlled largely by the stage with the smallest total probability of being replaced. A stage that is rarely replaced acts as a *bottleneck*: once patches enter this state, they remain there for many time steps, and so any transition that moves into this stage relaxes very slowly (Fig. 2A). As a result, in model (1) this stage accumulates the largest share of patches (however, this need not be the case in more complicated transition matrices). Consequently, adding or removing transitions between other, faster stages does little to change resilience unless those changes alter the exit rate of this bottleneck stage.

**Figure 2:**
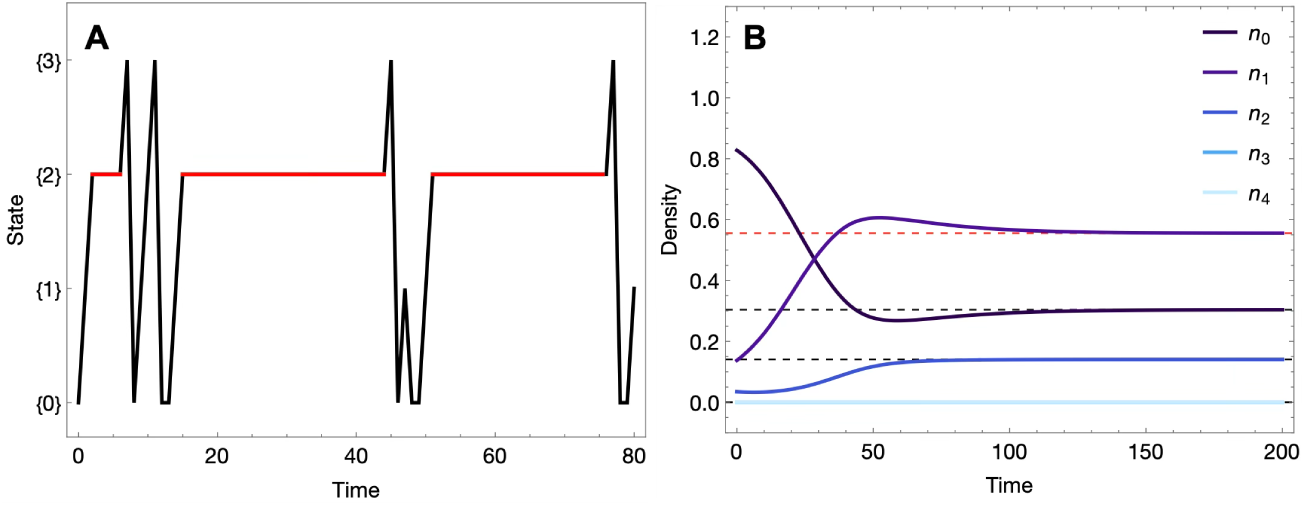
Conceptual diagram of bottleneck species in time. A) Time dynamics of Markov model in Eq. 1, highlighting bottleneck species (species 1 in this case) dominating the equilibrium abundance. B) Time dynamics from the continuous succession model in Eq. (5) with a perturbation from equilibrium as the initial condition. Equilibrium frequencies are represented by black dashed lines, with red dashed line representing bottleneck frequency.

### Time to reach the final stage

Thus far, resilience has referred to the rate at which the community approaches its stationary distribution. We now consider a different quantity: the expected time required for an individual patch to reach the terminal successional stage, or “climax” stage. In the context of succession, this quantity is useful as the prevalence of these stages often signifies the maturity of a successional community (Halpern 1988; Lebrija-Trejos et al. 2010). To this end, one can compute the “mean first passage time” (MFPT), or the expected number of species replacements during succession which must occur until the final stage is reached from a particular starting stage (Grinstead and Snell 2012).

To calculate this, denote *m_i_* as the time to reach the final state *S* from stage *i*. One can solve the MFPT *m_i_* by writing a system of recursive equations as:

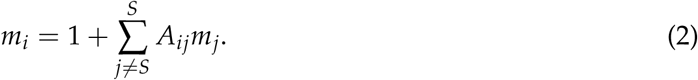

In other words, the time to go from stage *i* to stage *S* is one step plus the time it takes to reach stage *S* from any preceding stage. For our matrix, the system of equations is as follows:

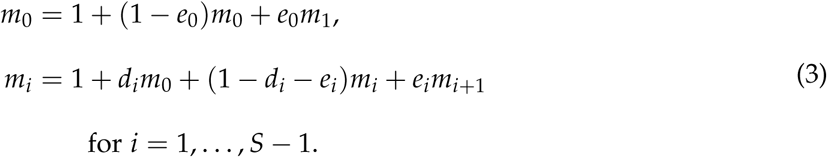

while the time to get from the final stage to itself is zero (*m_S_* = 0). Focusing on the majority of stages, one can rearrange Eq. (3) to get an implicit expression for the time it takes most states to reach the final stage:

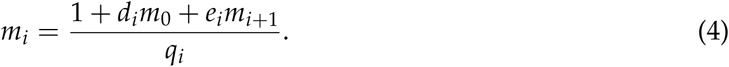

Note that in the denominator is the exit rate *q_i_* from stage *i*. Because MFPT is the sum of expected waiting times for all later stages, stages with low exit rates contribute disproportionately to total passage time. As the bottleneck stage has the smallest exit rate *q_i_*, it has the largest contribution to the MFPT, so increasing its exit rate reduces the dominant waiting-time component of succession.

As a result, perturbing the bottleneck lowers MFPT not only from that stage itself, but also from earlier and later stages whose trajectories spend time there (Fig. 3). Thus, the bottleneck stage controls the overall pace of succession because every successional pathway is delayed by its long residence time.

**Figure 3:**
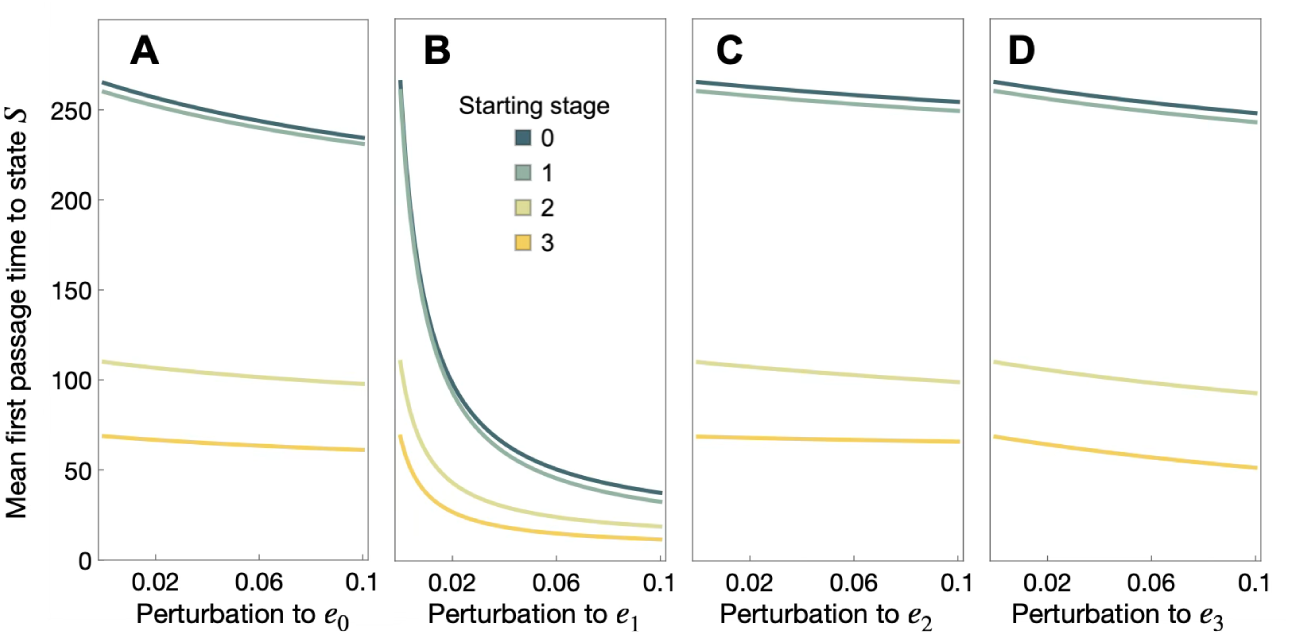
Mean first-passage time (MFPT) to the final successional stage from each starting stage in a five-species example of model (1). Panels A–D show the effect of increasing the colonization rate *e_i_* of species *i* = 0, 1, 2, 3, respectively. Stage 1 is the bottleneck stage, so increasing *e*_2_ (panel C) produces a substantially larger reduction in MFPT from all starting stages than equivalent perturbations to the other species. Parameters: (*e*_0_ = 0.2, *e*_1_ = 0.01, *e*_2_ = 0.4, *e*_3_ = 0.3, *d*_0_ = 0.1, *d*_1_ = 0.1, *d*_2_ = 0.1, *d*_3_ = 0.3)

### Density-dependent succession

A common criticism of Markov models of succession is that they treat replacement probabilities as fixed, and therefore neglect density dependence in species replacement (Usher 1981). In real successional communities, however, replacement rates often depend on which species are locally abundant and able to disperse into a patch. This raises the question: is the bottleneck principle merely a consequence of linear, density-independent replacements, or does it persist when replacement depends on the density of competitors? It is straightforward to show that the same qualitative intuition holds in a density-dependent model of succession. When self-replacement is strong as in many empirical successional systems, the stage with the lowest effective exit rate still dominates the slowest mode of return, indicating that bottlenecks are not simply an artifact of linear Markov assumptions but a more general consequence of asymmetries in persistence.

To address this, we study the continuous-time nonlinear model of succession analogue to Eq. (1) which is well-studied (Horn 1975; Miller and Allesina 2021) .Consider a guild of *S* species with abundances *N_j_*(*t*), *j* = 1, . . ., *S*. We use *j* to index the focal species and *i* to index source species contributing to its recruitment. Their dynamics are governed by

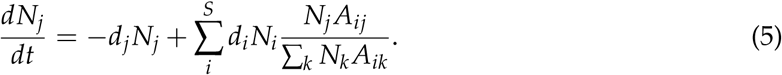

Here, *d_j_* represents the disturbance rate (or turnover rate) associated with state *j*, determining how quickly biomass turns over from that state. The term *A_ij_* denotes the transition probability from state *i* to state *j* upon disturbance, such that ∑*_j_ A_ij_* = 1. In words, the growth of state *i* is reduce at rate *d_j_N_j_* and is redistributed into other states according to the disturbance rates *d_j_* of their source states, weighted by the transition probabilities *A_ij_*. The total flow between states therefore depends both on how frequently each state is disturbed and on where individuals are likely to end up afterward.

The dominant eigenvalue of the Jacobian evaluated at an equilibrium *N*^∗^ determines the relaxation rate to equilibrium after perturbations, or resilience in continuous-time models (DeAngelis 1980; Pimm 1984):

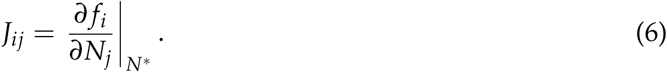

Unlike in the discrete, linear model, in the nonlinear model the rate that equilibrium is attained is limited not only by one species, but by the entire web of interacting species. However, it is straightforward to show that the bottleneck species that are unlikely to be replaced determine resilience most strongly.

Suppose that our matrix follows Eq. (1), meaning that most off-diagonal entries are zero. Moreover, since most successional matrices are dominated by the diagonal (Baasch et al. 2010; Tanner et al. 1996) (also see Fig. 1), the diagonal of the Jacobian is most relevant for determining the rate of return to equilibrium. The Jacobian of model (5) evaluated at an equilibrium *N*^∗^ (following (Horn 1975)):

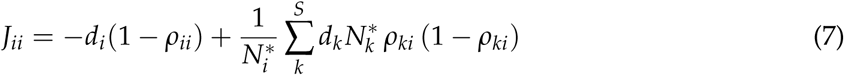

where the frequency of each species given their density in the community is defined as:

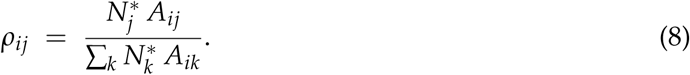

The first term, *ε_i_* = *d_i_*(1 − *ρ_ii_*), is the effective exit rate from stage *i*. Individuals are disturbed at rate *d_i_*, but only the fraction 1 − *ρ_ii_* transitions to another stage, while the remaining fraction *ρ_ii_* returns to the same stage. At the bottleneck, *ρ_ii_* ≈ 1, so the effective exit rate is small. The second term represents the contribution of recruitment into stage *i*.

We can relate the recruitment rate to the exit rate using the equilibrium condition from Eq. (5). At equilibrium, departures from stage *i* must be balanced by recruitment from all other stages to stage *i*:

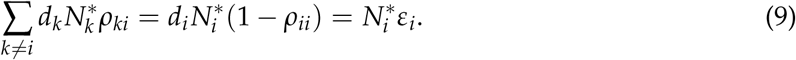

Thus, when the effective exit rate is small, the total recruitment flux into stage *i* from other stages must also be small. The corresponding part of the Jacobian contains these same recruitment fluxes, multiplied by 1 − *ρ_ki_*, a quantity between zero and one. It therefore cannot be larger than *ε_i_* and hence becomes small as *ε_i_* becomes small.

Consequently, when *ρ_ii_* ≈ 1, both the loss term and the recruitment term in *J_ii_* are small. The bottleneck therefore limits both the rate of exiting and entering its stage, slowing the overall pace of succession. Since resilience is determined by the dominant non-zero eigenvalue of the Jacobian, the bottleneck is therefore expected to largely determine community resilience.

## Discussion

We have introduced a simple and broadly useful principle of ecological succession which effectively reduces the complexity of studying resilience in communities. Although succession is typically viewed as the outcome of many interacting transitions among species, both return rates and passage times can often be well-approximated by a much simpler description in which a single bottleneck stage governs the dominant timescale of community dynamics. While classical intuition on succession suggests that the path to equilibrium is “rate limited” by slow species (Jakovac et al. 2021; Whittaker 1974), we have formalized this logic and extended a counter-intuitive corollary: increasing transition rates elsewhere in the network, or adding alternative successional pathways, has little effect on resilience unless those changes alter the bottleneck stage itself.

While our analysis is framed using Markov models of succession, the bottleneck principle does not depend on this formalism. Any process of community development characterized by stages with differing residence times will tend to be governed by its slowest transition. A “stubborn” intermediate species may therefore dominate the return to equilibrium even when succession does not follow from a mechanism not studied here (e.g. priority effects, intransitive competition, etc.). Transition matrices simply provide a convenient way of identifying bottlenecks, but the underlying principle is caused by slow-transitioning species slowing the development of the community towards a mature stage.

In practice, candidate bottlenecks can be identified directly from empirical transition matrices. Because bottleneck stages are both slow to exit and disproportionately affect recovery to equilibrium, they typically have low effective exit rates and/or large elements in the subdominant eigenvector of the transition matrix. The first quantity identifies stages with long residence times, while the second identifies those that contribute most strongly to *λ*_2_. Together, these provide a simple empirical diagnostic for identifying stages most likely to constrain successional development.

Arrested succession at a specific stages is often viewed as an alternative stable state (Young et al. 2001). Our results suggest another possibility; some apparently stable states may instead represent long-lived transients generated by exceptionally low exit rates. In such systems, a return to equilibrium remains possible, but occurs so slowly that the community appears effectively stationary over observational timescales. Distinguishing between true alternative stable states and bottleneck-driven transients may therefore require estimating transition rates and return timescales, rather than relying solely on the apparent persistence of a specific community state.

It would be interesting to study the life history strategies and traits associated with bottleneck species. For example, some grassland plants with poor dispersal have a tendency to arrest development as they can effectively halt the community at an alternative stable state (Cramer et al. 2008). Alternatively, some species of bamboo tend to exclude trees by employing fast growth in early stages to close the canopy and limit light for invaders (Zheng and Pacala 2024). Across systems, bottlenecks are therefore likely to arise not from any single life-history strategy, but from any trait combination that facilitates long residence times and suppresses replacement.

Many of the results here have strong implications for ecological management and restoration. Assuming that replacement probabilities remain approximately constant, improving the establishment or persistence of species other than the bottleneck will have relatively little influence on the overall rate of succession. Instead, restoration efforts should first identify the bottleneck stage and target interventions that increase its effective exit rate, either by facilitating replacement by later-successional species or by increasing the frequency of disturbances that promote its turnover. By identifying and relieving the principal bottleneck, managers can achieve accelerated community resilience.

## Conclusion

Despite the complexity of succession, we have shown that resilience can be assigned a simple biological interpretation through the concept of bottlenecks. Identifying bottleneck stages by understanding their life histories, traits, or behaviors may aid in ultimately enhancing resilience across a wide range of successional communities.

## Acknowledgements

The insightful comments and suggestions of Vincent Pan, Keila Stark, Jonas Wickman and Jonathan Levine substantially improved the clarity of the manuscript. This material is based upon work supported by the National Science Foundation Graduate Research Fellowship under Grant No. 2235783. This is W.K. Kellogg Biological Station contribution #XXXX.

## Statement of Authorship

N.R. conceived the study, performed the analyses, and wrote the manuscript.

## Data and Code Availability

Upon submission, data and code supporting this study will be deposited to a Zenodo repository containing the Mathematica scripts including empirical transition matrices used in the analyses.

